# obaDIA: one-step biological analysis pipeline for data-independent acquisition and other quantitative proteomics data

**DOI:** 10.1101/2020.05.28.121020

**Authors:** Jun Yan, Hongning Zhai, Ling Zhu, Sasha Sa, Xiaojun Ding

## Abstract

**Motivation:** Data mining and data quality evaluation are indispensable constituents of quantitative proteomics, but few integrated tools available.

**Results:** We introduced obaDIA, a one-step pipeline to generate visualizable and comprehensive results for quantitative proteomics data. obaDIA supports fragment-level, peptide-level and protein-level abundance matrices from DIA technique, as well as protein-level abundance matrices from other quantitative proteomic techniques. The result contains abundance matrix statistics, differential expression analysis, protein functional annotation and enrichment analysis. Additionally, enrichment strategies which use total proteins or expressed proteins as background are optional, and HTML based interactive visualization for differentially expressed proteins in the KEGG pathway is offered, which helps biological significance mining. In short, obaDIA is an automatic tool for bioinformatics analysis for quantitative proteomics.

**Availability:** obaDIA is freely available from https://github.com/yjthu/obaDIA.git.

## 1 Introduction

Data-independent acquisition (DIA) proteomics technology is getting more popularity and recognition recently. Compared with data-dependent acquisition (DDA) strategy, DIA avoids the topN acquisition randomness and has better reproducibility, and more suitable for large cohort research. Many tools have been developed to identification and quantification DIA data, such as OpenSWATH, Spectronaut, and Skyline (Zhang, *et al*., 2020). However, limited tool is specifically designed for one-step, fast and comprehensive downstream bioinformatics analysis of DIA data.

Some biological data analysis frameworks provide tools for downstream biological analysis of quantitative proteomics data and personalized workflow construction. For example, R/Bioconductor(Gentleman, et al., 2004) provides MSstats(Choi, et al., 2014) and DEqMS(Zhu, et al., 2020) for expression analysis, topGO(Alexa and Rahnenführer, 2009) and clusterProfile(Yu, et al., 2012) for function analysis; Bioconda(Gruning, et al., 2018) or Galaxy(Jalili, et al., 2020) offers BLAST(Altschul, et al., 1990), HMMER(Potter, et al., 2018) and DIAMOND(Buchfink, et al., 2015) for functional annotation, as well as KOBAS(Mao, et al., 2005) for enrichment analysis. Other independent software, such as mapDIA(Teo, et al., 2015), Perseus(Tyanova and Cox, 2018) and Automated-workflow-composition (Palmblad, et al., 2019), also contribute to specific downstream proteomics data mining. But an integrate one-step software is still necessary for proteomic community, considering some users prefer to obtain the standard downstream analysis results quickly. Here, we introduce obaDIA, an all-in-one pipeline which generates comprehensive and reproducible biological analysis results from DIA and other quantitative proteomics data automatically.

## 2 Application Description

### 2.1 Workflow of obaDIA

obaDIA takes two standard format files (TSV format abundance matrix and FASTA format protein sequences) as inputs and automatically performs abundance imputation, normalization, differential expression, functional annotation and enrichment analysis to produce results. It supports fragment-level, peptide-level and protein-level abundance matrices from DIA technology, and protein-level abundance matrices from other quantitative proteomics technologies. The obaDIA workflow has three core modules. The “AB & DE” module is used for abundance matrix (AB) process and batch differential expression (DE) analysis. The “Annotation” module carries out detailed functional annotation and classification for expressed proteins against SWISS-PROT, PFAM, GO, KEGG, eggNOG and signalP databases. The “Enrichment” module receives the output files from “AB & DE” and “Annotation” as inputs and performs functional enrichment analysis of GO, KEGG, PFAM and Rectome databases for differentially expressed proteins from all predefined comparisons (Figure 1).

**Figure 1.**
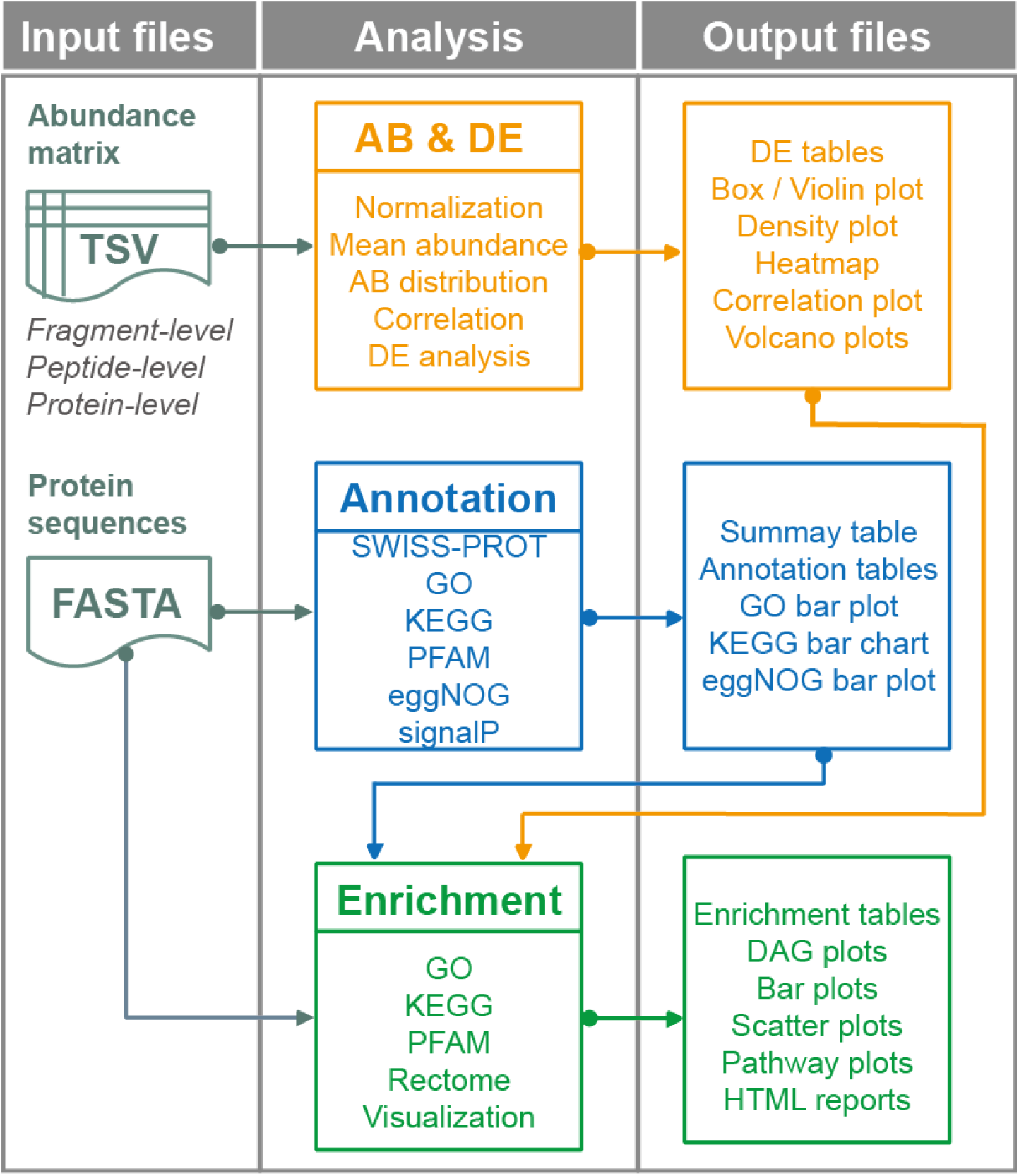
Diagram of the workflow of obaDIA.

### 2.2 Features of obaDIA

The obaDIA pipeline is a one-step solution to perform multiple downstream biological analysis and generate comprehensive statistical charts. Although it is all-in-one, each module can be run separately, facilitating integration into personalized workflows. Specially, an HTML formatted report can be generated, and the information of expressed proteins can be displayed in a KEGG pathway interactively, which facilitates biological information mining. obaDIA takes protein-level abundance matrices as inputs for both DIA and other techniques, which ensures its wide application in the proteomics community; it also takes fragment-level or peptide-level abundance matrices as inputs for DIA data, which produces more accurate difference analysis results by using the mapDIA algorithm(Teo, et al., 2015). In addition, obaDIA provides two enrichment strategies for users, including the traditional enrichment strategy which uses total proteins as background, and an alternative strategy which uses expressed proteins in the experiment as background. Finally, in fast analysis mode, you can get all the main analysis results in a few minutes.

## 3 Implementation and Results

The main program of obaDIA was developed in Perl, and incorporates R, Python, and Bash scripts. All third-party tools and dependencies are listed in Table 1. We also provide a docker image, which greatly simplifies the installation of third-party dependencies, as well as a GUI for Galaxy users. The obaDIA pipeline has been tested on Linux platform CentOS release 7.0 and should work on similar Linux/Unix systems which support Bash command line. The source code, wrapped dependencies and user manual of obaDIA can be obtained from https://github.com/yjthu/obaDIA.git.

**Table 1.** Integrated tools and dependencies in obaDIA.

| Module | Dependencies | References | Wrapped |
| --- | --- | --- | --- |
| AB & DE* | R packages: corrplot, pheatmap,<br>PerformanceAnalytics |  | no |
|  | mapDIA | (Teo, *et al.*, 2015) | yes |
| Annotation | Trinotate-v3.1.0-pro (a modified<br>version) | (Bryant, *et al.*,<br>2017) | yes |
|  | sqlite3 |  | system |
|  | DIAMOND | (Buchfink, *et al.*,<br>2015) | yes |
|  | hmmScan & hmmpress | (Potter, *et al.*, 2018) | yes |
|  | signalP 4.1 | (Petersen, *et al.*,<br>2011) | no |
|  | Perl5 libraries: GO::Parser, DBI,<br>DBD::SQLite |  | no |
| Enrichment | KOBAS 3.0 | (Mao, *et al.*, 2005) | no |
|  | R packages: GO.db, goseq, goTools,<br>topGO, Rgraphviz |  | no |
|  | Python2 libraries: eventlet,<br>bs4,django==1.8.3 |  | no |
| Basic | R packages: reshape2, ggplot2,<br>optparse |  | system |
|  | Perl5 |  | system |
|  | Python2 |  | system |

\*AB: abundance analysis, DE: differential expression analysis

The obaDIA pipeline has been tested using a published demo data(Teo, et al., 2015) and an unpublished complete DIA data. All input and output files from the published demo data set are wrapped with the obaDIA release, and the main results are also shown and described in Supplementary Figure 2-6. On our workstation (Table 2), the obaDIA workflow can be completed within two hours in standard mode and within five minutes in fast mode. In terms of enrichment analysis results, we compared the results of two built-in enrichment strategies based on total proteins and expressed proteins, respectively. The P-value and order of the enrichment items are different (Figure 7); users should choose the enrichment method suitable for their own data according to the biological significance of the results. In summary, obaDIA provides an integrated one-step biological analysis solution, which may facilitate standardized biological analysis for proteomics data.

**Table 2.** Time consumption of obaDIA workflow in test data sets.

| Test data description | Step | Running time | Running time |
| --- | --- | --- | --- |
|  |  | (minutes) * | (minutes) |
|  |  | *default mode* | *fast mode* |
| Data set 1: | AB & DE | 1 | 1 |
| Species: Human; | Annotation | 13 | 1 |
| Six groups and each group | Enrichment | 59 | 2 |
| with three biological |  |  |  |
| replicates; | Total | 73 | 3 |
| 566 proteins, 5 comparisons |  |  |  |
| Data set 2: | AB & DE | 1 | 1 |
| Species: Mouse; | Annotation | 46 | 2 |
| Four groups and each group | Enrichment | 71 | 1 |
| with three biological |  |  |  |
| replicates; | Total | 118 | 4 |
| 4493 proteins, 2 |  |  |  |
| comparisons |  |  |  |

\* The main factor affecting the running time in different systems is CPU. The CPU we used for testing is: Intel(R) Xeon(R) CPU E5-2687W v3 @ 3.10GHz. We used 40 threads for hmmscan in order to accelerate annotation. One thread was used for the other steps. If the time is less than one minute, count as one minute.

**Figure 2.**
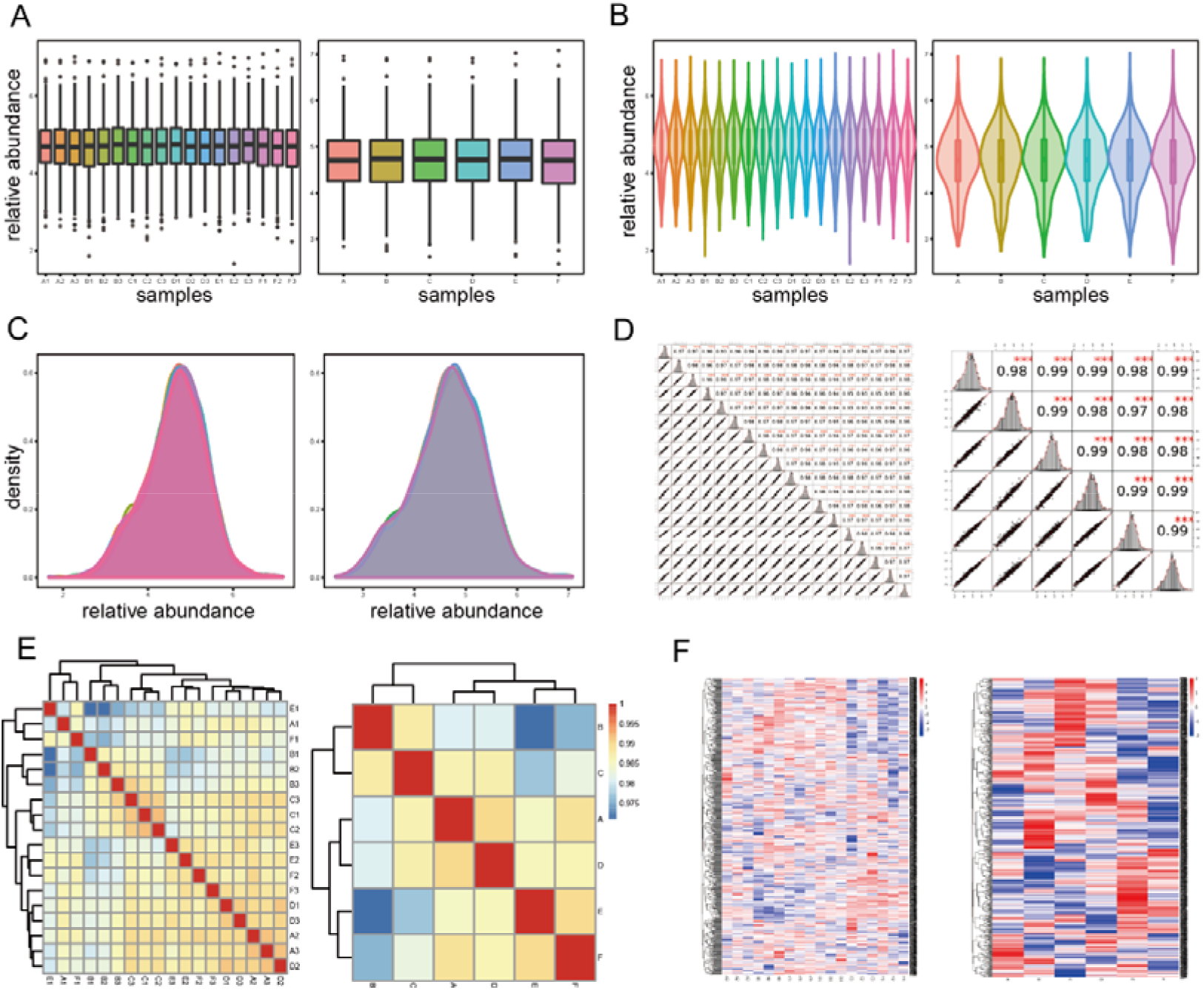
Protein abundance analysis results. The raw abundance matrix is normalized to generate a relative abundance matrix and a group mean abundance matrix, then they are scaled to log_10_(*value*+1) for plotting. (A) Box plot of relative abundance for each sample (left) and group (right). (B) Violin plot of relative abundance for each sample (left) and group (right). (C) Density plot of relative abundance for each sample (left) and group (right). (D) Correlation matrix for each comparison of samples (left) and groups (right). Left-bottom shows scatter plots, right-top shows Pearson correlation coefficients (PCC), and diagonal shows histograms of of the relative abundance. An asterisk indicates statistical significance. (E) Correlation heatmap for each comparison of samples (left) and groups (right). Red color indicates higher PCC while blue indicates lower. (F) Relative abundance heatmap for each sample (left) and each group (right). Red color indicates higher abundance while blue indicates lower.

**Figure 3.**
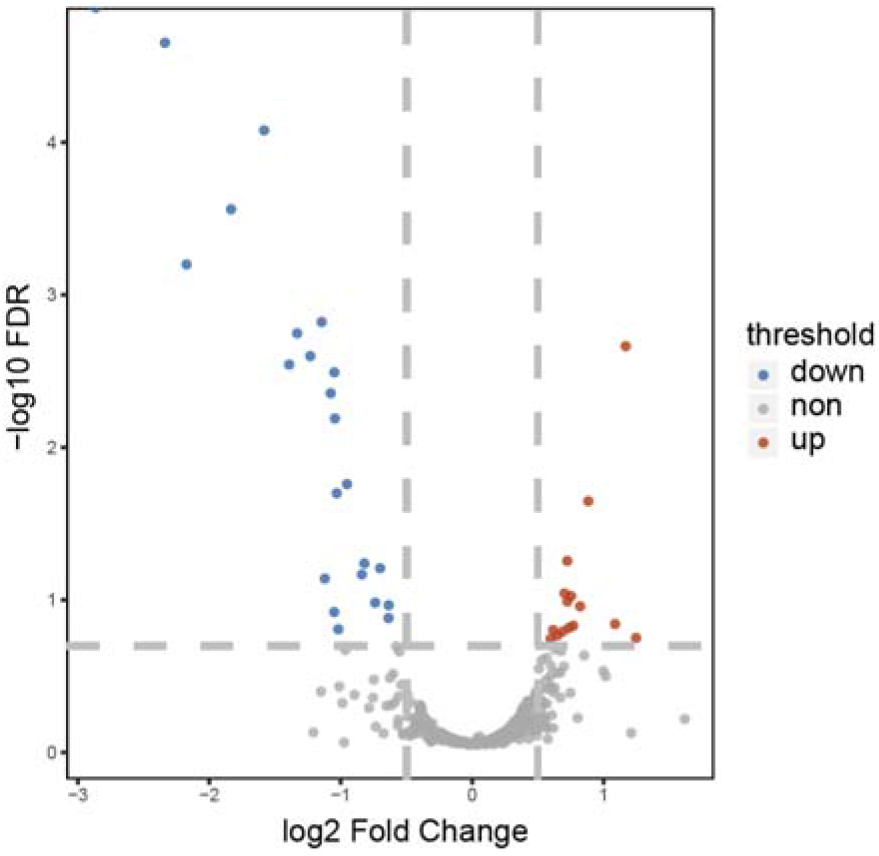
Volcano plot for differential expressed proteins filtering. Red, blue and grey dots represent up, down expressed and non-DE proteions respectively. A greater absolute value on the X-axis indicates a greater expression difference, while a higher value on the Y-axis indicates a more significant expression difference between the two samples.

**Figure 4.**
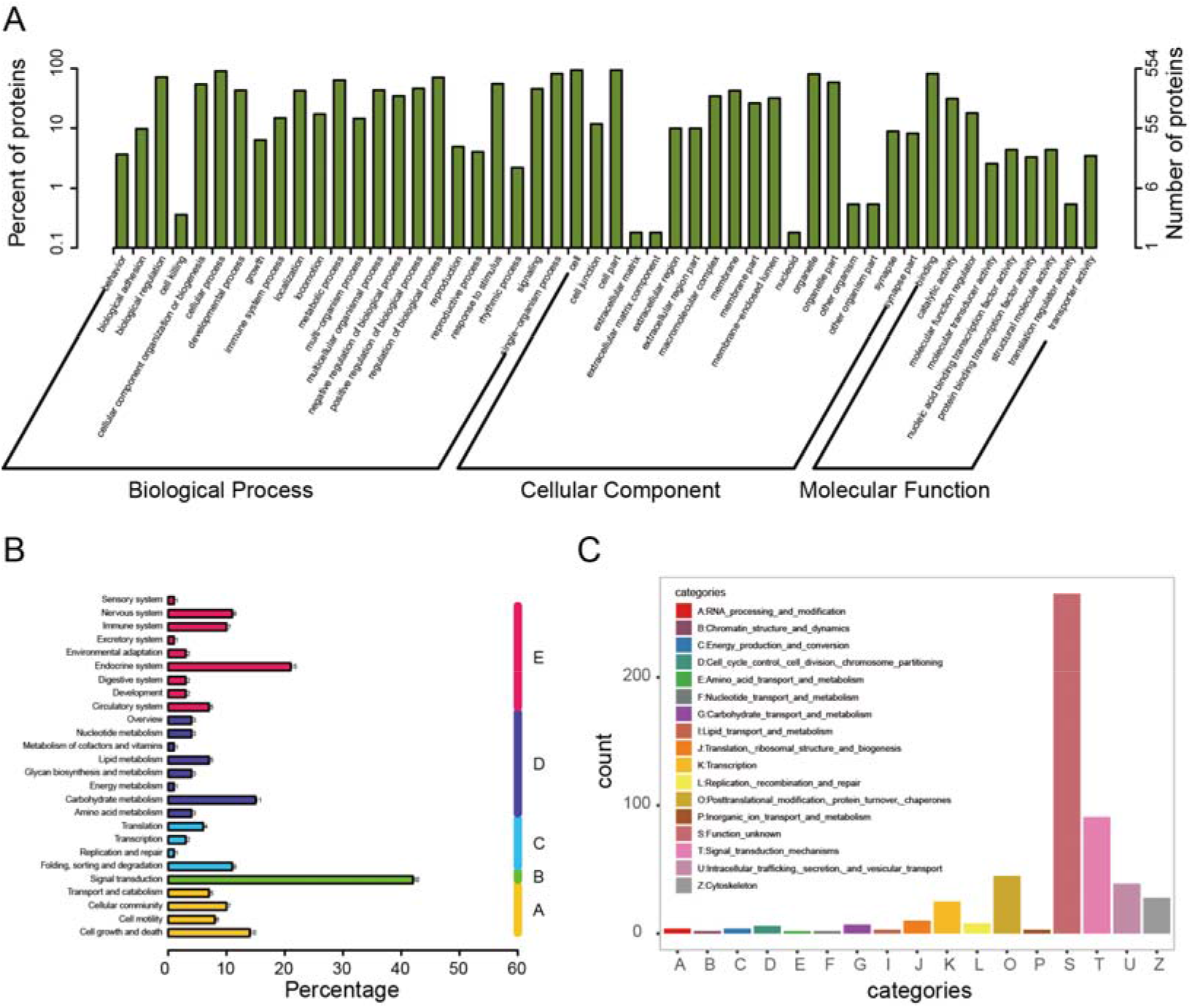
GO, KEGG and eggNOG classification of protein functional annotation results. (A) Bar plot of GO classification. Top 50 second-level GO terms are ploted based on the first-level GO categories: biological process, cellular component and molecular function. The number and percentage of proteins in each GO term(including its child terms) are shown. (B) Bar chart of KEGG classification. All KEGG terms are assigned to five main categories: A (Cellular Processes, B (Environmental Information Processing, C (Genetic Information Processing), D (Metabolism), and E (Organismal Systems). (B) Bar plot of eggNOG classification. All eggNOG terms are assigned into categories A-Z.

**Figure 5.**
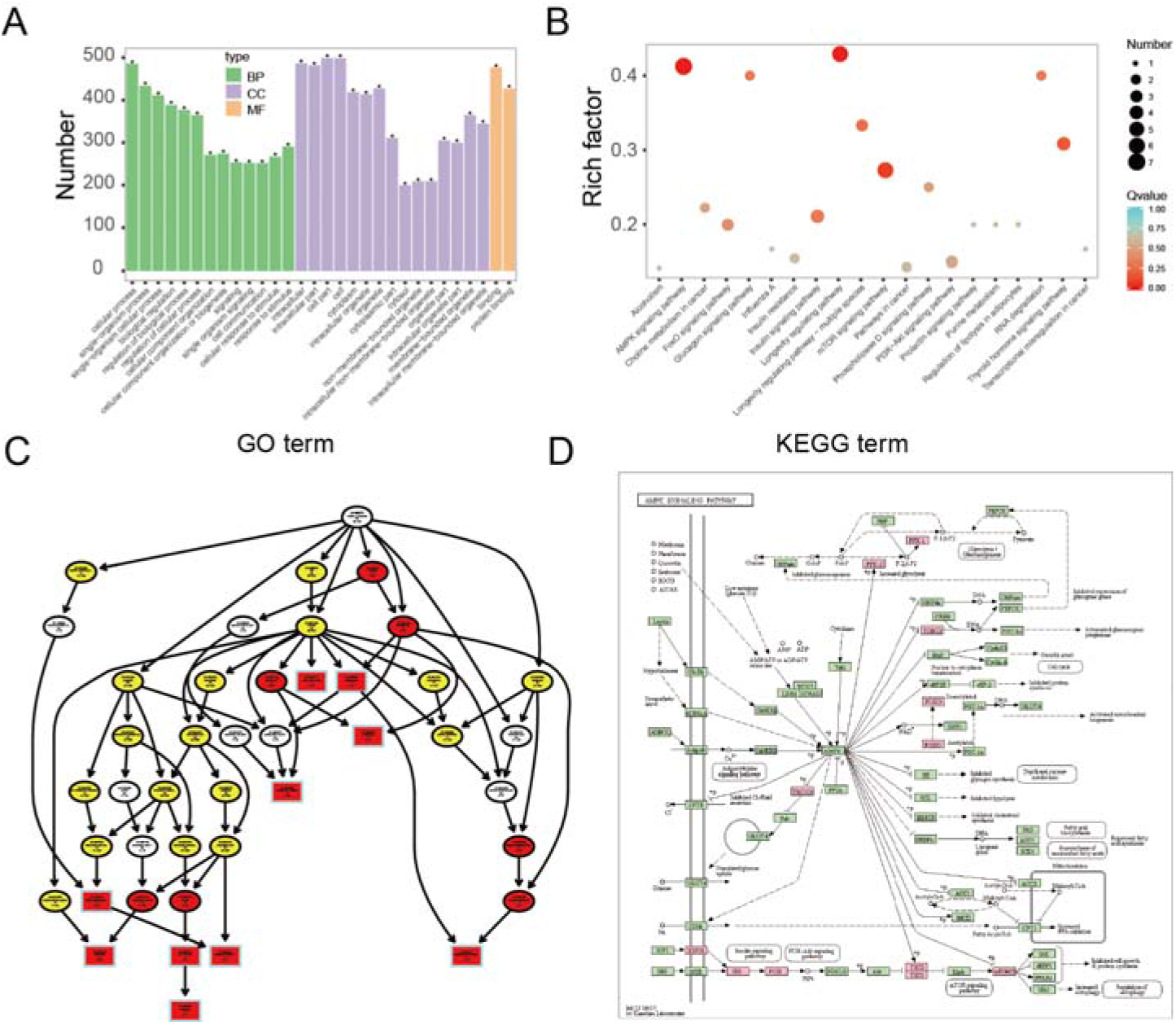
Representative results of functional enrichment analysis. (A) Bar plot of GO enrichment. Top 30 GO terms are ploted based on three categories: BP (biological process), CC (cellular component) and MF (molecular function). An asterisk indicates statistical significance. (B) Scatter plot of KEGG enrichment. Rich factor is the ratio between the input number and the total number of proteins in the same pathway. (C) DAG (Directed Acyclic Graph) plot of GO encrichment. Branches represent the inclusion relationship, and the function scope defined from top to bottom is getting smaller and smaller. Generally, the top 10 most enriched GO terms are displayed with rectangles while others with ellipticals, and different degrees of enrichment are shown in red, yellow and white, respectively. GO term identifier, description, number of proteins in the term are shown in each node. (D) A pathway plot obtained through KEGG API. Expressed proteins in the pathway nodes are shown in pink.

**Figure 6.**
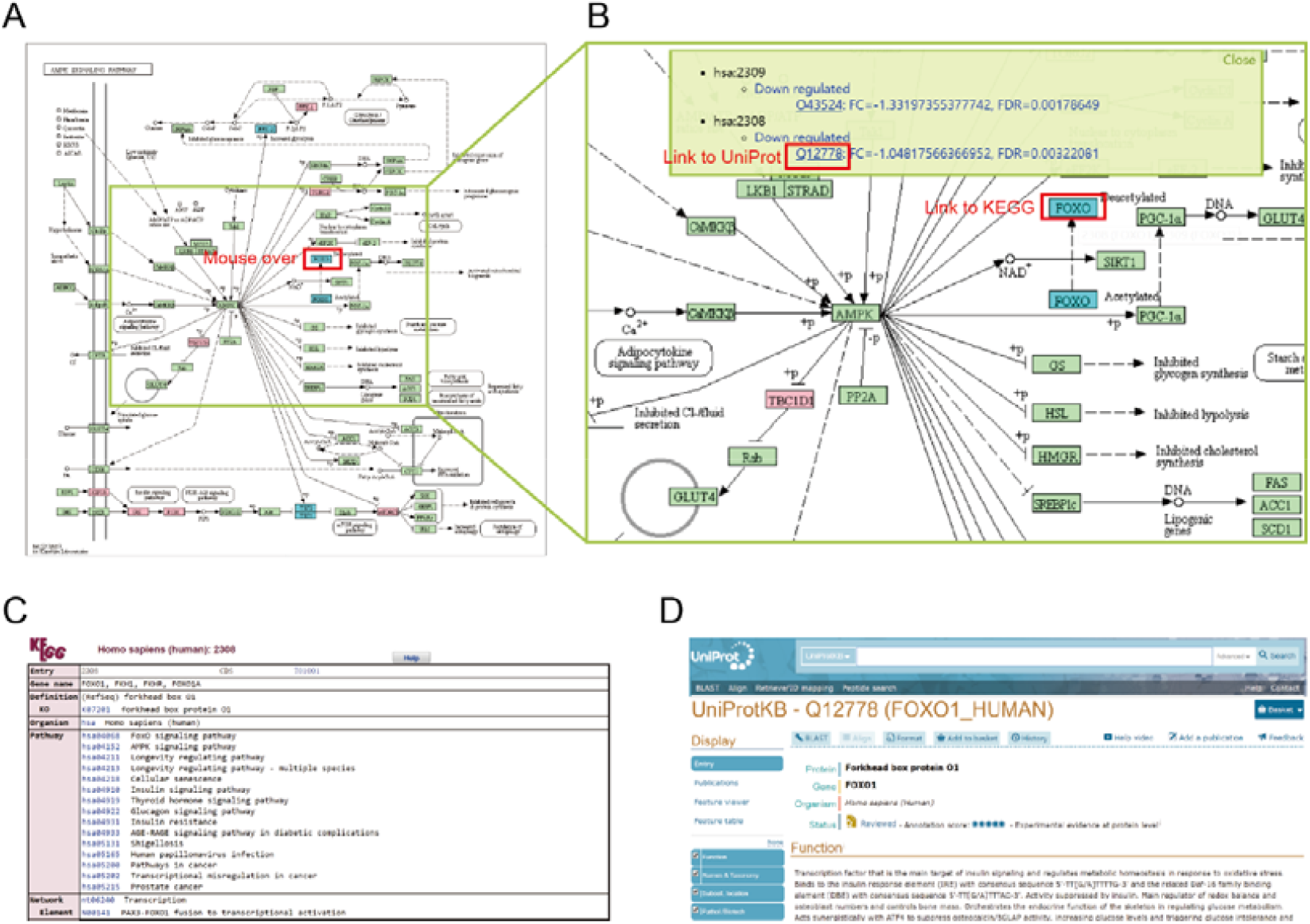
KEGG pathway visualization. (A) AMPK signaling pathway plot obtained through KEGG API. Expressed proteins are shown in pink and differentially expressed proteins are shown in red (upregulation, there is no upregulated protein in this example) or blue (down- regulation). (B) Local amplification of the AMPK signaling pathway plot. A display box pops up upon mouse-over of a down regulated protein *FOXO*, which shows the protein UniProt identifier, fold change and FDR values of the differential expressed analysis. Clicking on a protein name node in the pathway will link to the KEGG website (C), and clicking on the protein UniProt identifier in the display box will link to the UniProtKB website (D).

**Figure 7.**
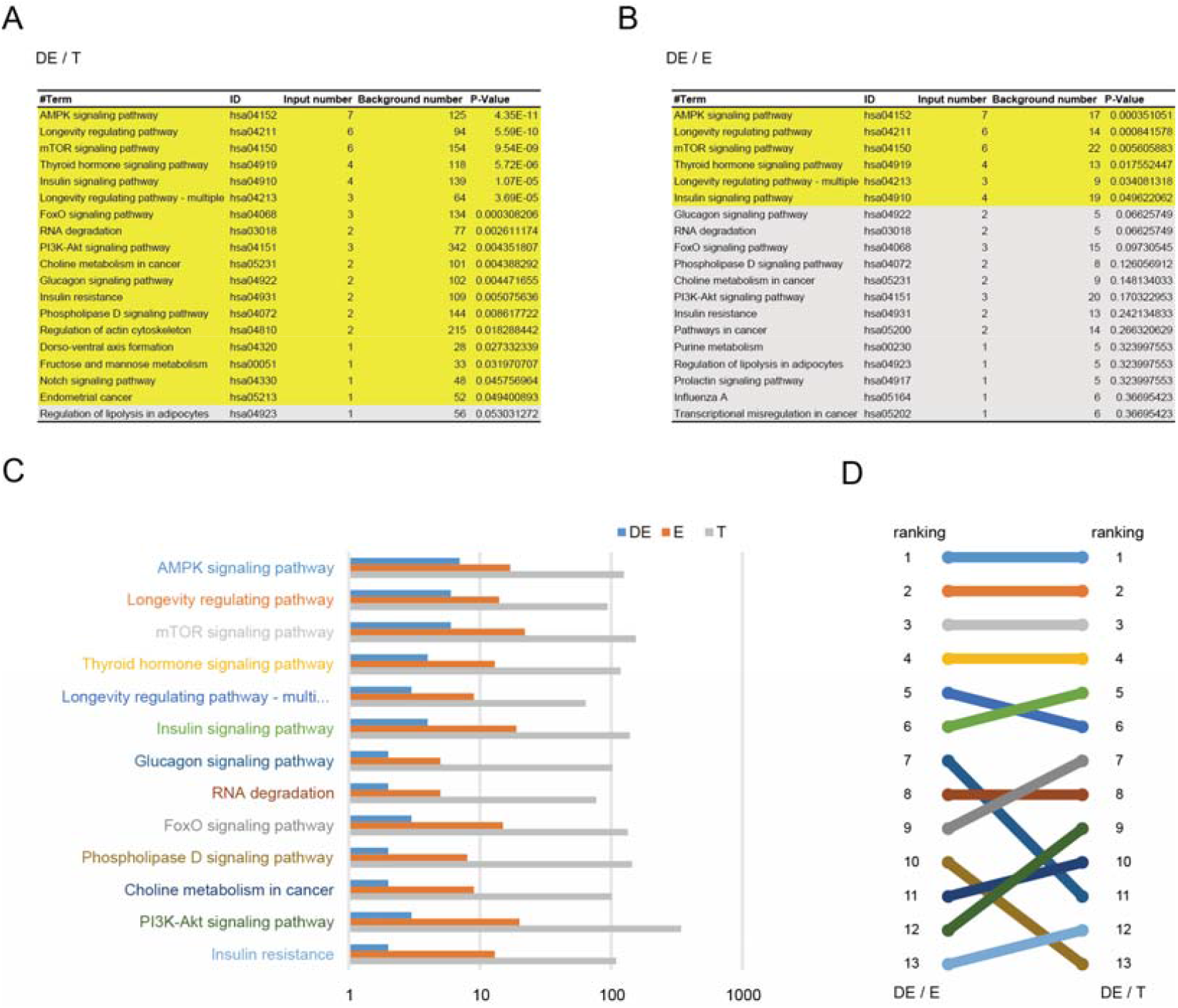
Comaprison of two enrichment methods. (A) Snapshot of KEGG enrichment results table used total (*T*) proteins in each pathway as backgroud; yellow highlights indicate the statistically significant (*P* < 0.05) terms. (B) Snapshot of KEGG enrichment results table using expressed (*E*) proteins in each pathway as backgroud; yellow highlights indicate the statistically significant(*P* < 0.05) terms. (C) Bar chart shows numbers of differentially expressed (*DE*), expressed (*E*) and total (*T*) proteins in each pathway. (D) Comaprison of the ranking of enriched KEGG terms using *E* as backgroud (*DE/E*) and using *T* as backgroud (*DE/T*) in the enrichment analysis results.

## Funding

Funding information is not applicable.

## Conflict of Interest

none declared.

## Notes

### Competing Interest Statement

The authors have declared no competing interest.

### Summary of Updates

A rectified version of the draft

## References

Alexa, A. and Rahnenführer, J. Gene set enrichment analysis with topGO. Bioconductor Improv 2009;27.

Altschul, S.F., et al. Basic local alignment search tool. J Mol Biol 1990;215(3):403–410.

Bryant, D.M., et al. A Tissue-Mapped Axolotl De Novo Transcriptome Enables Identification of Limb Regeneration Factors. Cell Rep 2017;18(3):762–776.

Buchfink, B., Xie, C. and Huson, D.H. Fast and sensitive protein alignment using DIAMOND. Nat Methods 2015;12(1):59–60.

Choi, M., et al. MSstats: an R package for statistical analysis of quantitative mass spectrometry-based proteomic experiments. Bioinformatics 2014;30(17):2524–2526.

Gentleman, R.C., et al. Bioconductor: open software development for computational biology and bioinformatics. Genome Biol 2004;5(10):R80.

Gruning, B., et al. Bioconda: sustainable and comprehensive software distribution for the life sciences. Nat Methods 2018;15(7):475–476.

Jalili, V., et al. The Galaxy platform for accessible, reproducible and collaborative biomedical analyses: 2020 update. Nucleic Acids Res 2020;48(14):8205–8207.

Mao, X., et al. Automated genome annotation and pathway identification using the KEGG Orthology (KO) as a controlled vocabulary. Bioinformatics 2005;21(19):3787–3793.

Palmblad, M., et al. Automated workflow composition in mass spectrometry-based proteomics. Bioinformatics 2019;35(4):656–664.

Petersen, T.N., et al. SignalP 4.0: discriminating signal peptides from transmembrane regions. Nat Methods 2011;8(10):785–786.

Potter, S.C., et al. HMMER web server: 2018 update. Nucleic Acids Res 2018;46(W1):W200–W204.

Teo, G., et al. mapDIA: Preprocessing and statistical analysis of quantitative proteomics data from data independent acquisition mass spectrometry. J Proteomics 2015;129:108–120.

Tyanova, S. and Cox, J. Perseus: A Bioinformatics Platform for Integrative Analysis of Proteomics Data in Cancer Research. Methods Mol Biol 2018;1711:133–148.

Yu, G., et al. clusterProfiler: an R package for comparing biological themes among gene clusters. OMICS 2012;16(5):284–287.

Zhang, F., et al. Data-Independent Acquisition Mass Spectrometry-Based Proteomics and Software Tools: A Glimpse in 2020. Proteomics 2020:e1900276.

Zhu, Y., et al. DEqMS: A Method for Accurate Variance Estimation in Differential Protein Expression Analysis. Mol Cell Proteomics 2020;19(6):1047–1057.

